# Mechanochemical Patterning Localizes the Organizer of a Luminal Epithelium

**DOI:** 10.1101/2024.10.29.620841

**Authors:** Sera Lotte Weevers, Alistair D. Falconer, Moritz Mercker, Hajar Sadeghi, Jaroslav Ferenc, Albrecht Ott, Dietmar B. Oelz, Anna Marciniak-Czochra, Charisios D. Tsiairis

## Abstract

The spontaneous emergence of tissue patterns is often attributed to biochemical reaction-diffusion systems. In *Hydra* tissue regeneration, the formation of a Wnt signaling center serves as a well-known example of such a process. However, despite extensive research, a strictly biochemical mechanism for self-organization in *Hydra* remains elusive. In this study, we investigated mechanical stimuli and identified a positive feedback loop between Wnt signaling and tissue stretching. We developed a mathematical model of mechanochemical pattern formation in a closed elastic shell, representing regenerating *Hydra* epithelial spheroids. Our model explains how mechanical forces drive axis formation and predicts the organizer’s location under various perturbations, providing a more comprehensive understanding of the spatial dynamics involved. Validation by partially confining regenerating tissues showed that the organizer indeed forms in regions with the greatest stretching potential. This work highlights a novel mechanochemical mechanism for luminal epithelium patterning, suggesting that mechanical forces, in addition to biochemical signals, play a crucial role in tissue regeneration and axis specification. Our findings offer broader implications for the role of mechanical forces in tissue organization in various biological systems, opening new avenues for investigating mechanochemical feedback in development and regeneration.

## INTRODUCTION

The role of mechanical signals, alongside biochemical cues, is increasingly recognized as crucial in determining cell fates and tissue patterning (Goodwin and Nelson 2021; Mammoto and Ingber 2010). In closed epithelial lumens, which are a recurring theme in morphogenesis, specific mechanical stimuli can arise from pressure exerted by intraluminal fluid (Chan and Hiiragi 2020; Chugh, et al. 2022; Hannezo and Heisenberg 2019). Extreme deformations of epithelial sheets have been documented as active superelasticity, and can be traced to the cells dynamic cytoskeletal organization (Latorre, et al. 2018). Regenerating fragments of *Hydra* tissue, which fold into closed spheres, provide a valuable model to study these phenomena. During regeneration in *Hydra*, an organism with ancient evolutionary origins, key cell fate decisions are governed by mechanisms conserved across metazoans (Holstein 2022).

The freshwater cnidarian *Hydra* has a simple tubular body structure, with a mouth and tentacles at the oral end of its main axis, and a foot at the aboral end, which anchors the organism to solid surfaces (Vogg, et al. 2019). Tissue fragments from the body of *Hydra*, consisting of two epithelial layers—the epidermis and gastrodermis—form hollow spheroids during regeneration (Fütterer, et al. 2003). Successful regeneration into complete, small animals hinges on forming a new organizing center, marked by high Wnt ligand expression. The spontaneous emergence of these centers has been attributed to a reaction-diffusion mechanism (Gierer and Meinhardt 1972), governed by the principle of local activation and long-range inhibition (LALI) (Meinhardt and Gierer 2000; Oster and Murray 1989; Veerman, et al. 2021). This mechanism involves specific nonlinear interactions between two morphogens with markedly different diffusion rates. In this context, Wnt ligands are seen as secreted factors that positively regulate their own expression and act as activators (Hobmayer, et al. 2000; Nakamura, et al. 2011). However, despite significant research, the relevant reaction-diffusion pair of morphogens, and in particular the long-range inhibitor, has yet to be identified (Wang, et al. 2023).

Recent theoretical studies have shown that mechanical forces and stimuli, in place of biochemical signals, can also generate robust patterns (Hiscock and Megason 2015). The role of mechanical stimuli as an alternative to biochemical signals has been explored in *Hydra* regeneration (Braun and Keren 2018; Wang, et al. 2023). Analyses linking morphogen production to mechanical properties, such as tissue curvature, suggest that mechanical factors can drive the emergence of new organizers. A positive feedback loop between tissue stretch and *Wnt* expression has been anticipated as a driver of head formation in *Hydra*, though the details of this self-organization remain unclear (Mercker, et al. 2015; Veerman, et al. 2021). Indeed, subsequent research has shown that mechanical stretching of *Hydra* spheroids is essential for organizer formation during regeneration (Ferenc, et al. 2021; Soriano, et al. 2009). These spheroids undergo cycles of osmotically driven inflation and deflation events, mechanically stretching the epithelial layers. Mechanical oscillations have been examined for their role in tissue patterning and cell fate (Kruse and Riveline 2011). In *Hydra, Wnt3* gene expression levels have been shown to correlate directly with the degree of tissue stretch. Moreover, when mechanical oscillations are prevented, regeneration failure can be rescued by forced *Wnt3* expression (Ferenc, et al, 2021). This clear link between mechanical stimuli and the Wnt signaling pathway suggests that tissue patterning in *Hydra* is driven by a mechanochemical mechanism.

## RESULTS

### Wnt signaling facilitates tissue stretching

In this study, we further explored the connection between Wnt signaling and mechanical stimulation during *Hydra* regeneration. Our primary question was whether Wnt signaling influences the mechanical behavior of regenerating *Hydra* spheroids. Using a transgenic line that overexpresses *Wnt3* in the epidermis (Nakamura, et al. 2011), we observed that these spheroids inflated to significantly larger sizes before deflating, although their inflation rate remained unaffected (Fig. 1A-D, Movies S1-2). To confirm the role of canonical Wnt signaling, we directly manipulated the stability of β-catenin using chemical methods. When regenerating spheroids were treated with 5 µM alsterpaullone—a small molecule that inhibits β-catenin degradation—Wnt signaling was activated even in the absence of Wnt ligands (Leost, et al. 2000). Under these conditions, the spheroids continued to inflate to extreme sizes (Fig. 1E-F, Fig. S1, Movies S3-4). This suggests that elevated Wnt signaling enhances tissue’s ability to stretch. This effect could be due to β-catenin’s role in stabilizing cell-cell junctions (Valenta, et al. 2012), which strengthens the tissue’s ability to withstand force, or due to increased cell elasticity from Wnt-induced EMT (Gonzalez and Medici 2014). Our findings reveal a positive feedback loop between Wnt signaling and mechanical tissue stretching: while Wnt signaling promotes greater tissue stretch, prior research has shown that stretching, in turn, activates *Wnt3* expression (Ferenc, et al. 2021).

**Figure 1:**
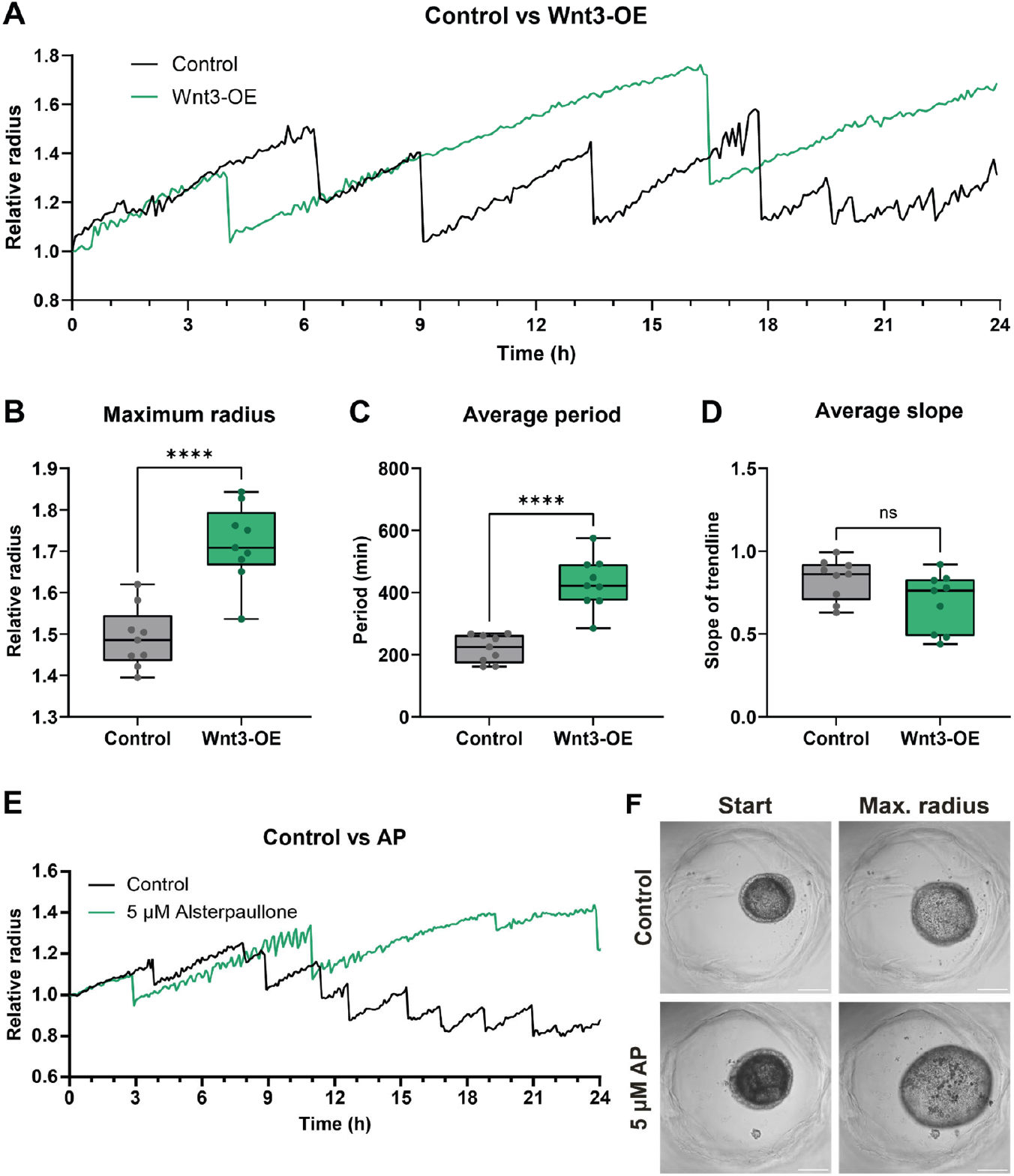
Wnt signaling enhances tissue stretching during *Hydra* spheroid regeneration. (**A**) Representative trajectories showing the size of regenerating, wild-type and *Wnt3* overexpressing spheroids over the first 24 hours of regeneration. Quantification of maximum radius (**B**), average period (**C**) and average slope (**D**) of the initial oscillations from regenerating, wild-type and *Wnt3* overexpressing spheroids (for both conditions, n = 9). (**E**) Representative trajectories showing the size of regenerating spheroids grown in *Hydra* medium supplemented with 0.2% DMSO or 5 μM alsterpaullone (AP) during the first 24h of regeneration. (**F**) Snapshots of the regenerating spheroids in (E). Shown are the initial time points and time points at which the spheroids have reached their peak size. Data was analyzed using an unpaired, two-tailed t-test. ^****^*P* < 0.0001. Scale bars, 200 μm.

### Mechanochemical mathematical model

Importantly, the discovered mechanochemical auto-activation of Wnt signaling surpasses the conventional biochemical short-range positive feedback loop postulated by the Gierer-Meinhardt model (Gierer and Meinhardt 1972; Nakamura, et al. 2011). The positive feedback loop between stretch and Wnt signaling causes excessive stretching in certain cells exposed to Wnt signaling, while the rest of the tissue remains relatively undisturbed. As a consequence, when some tissue regions stretch more, other areas compensate by stretching less, adjusting to the total volume of the lumen. Therefore, unlike classical reaction-diffusion models for pattern formation, our model does not require a rapidly diffusing inhibitor as this mechanism provides a long-range inhibitory effect in a natural manner.

In order to assess the patterning capacity of this regulatory mechanism, we constructed a mathematical model that represents *Hydra* spheroids as inflated elastic thin shells that undergo periodic inflation and deflation (supplementary text). Our model couples stress balance with the reaction-diffusion kinetics of a morphogen (Fig. 2A-B). The feedback loop is modeled by having the elastic modulus decrease with increasing morphogen concentration, while the morphogen production rate increases with tissue strain (Fig. 2C, supplementary text). Simulations of this model exhibit spontaneous pattern formation similar to that observed in regenerating *Hydra* spheroids. Specifically, a spherical shell with initially random, low levels of morphogen expression (Fig. 2D) can develop localized high morphogen expression and tissue deformation (Fig. 2E-F). Typically, a single organizer emerges in regions experiencing progressive stretching (Movie S5). The model’s ability to break symmetry critically depends on the degree of stretching and the degradation rate of the morphogen. Insufficient stretching, caused by reduced water influx, and rapid morphogen turnover due to high degradation levels prevent the positive feedback loop from amplifying initial variability (Fig. S2). Interestingly, previous observations show a corresponding gradual decline in spheroid regenerative capacity as the osmotic environment approaches isotonic conditions (Ferenc, et al. 2021). Moreover, the significant increase in spheroid volume observed under Wnt stimulation (Fig. 1A) is reproduced in our simulations when the equations include a constant source of additional morphogen expression (Fig. 2G). Notably, the behavior seen with experimental *Wnt3* overexpression naturally arises in the simulations without any prior adjustments aimed at achieving this result. Thus, our mathematical simulations align with and help clarify earlier experimental findings.

**Figure 2:**
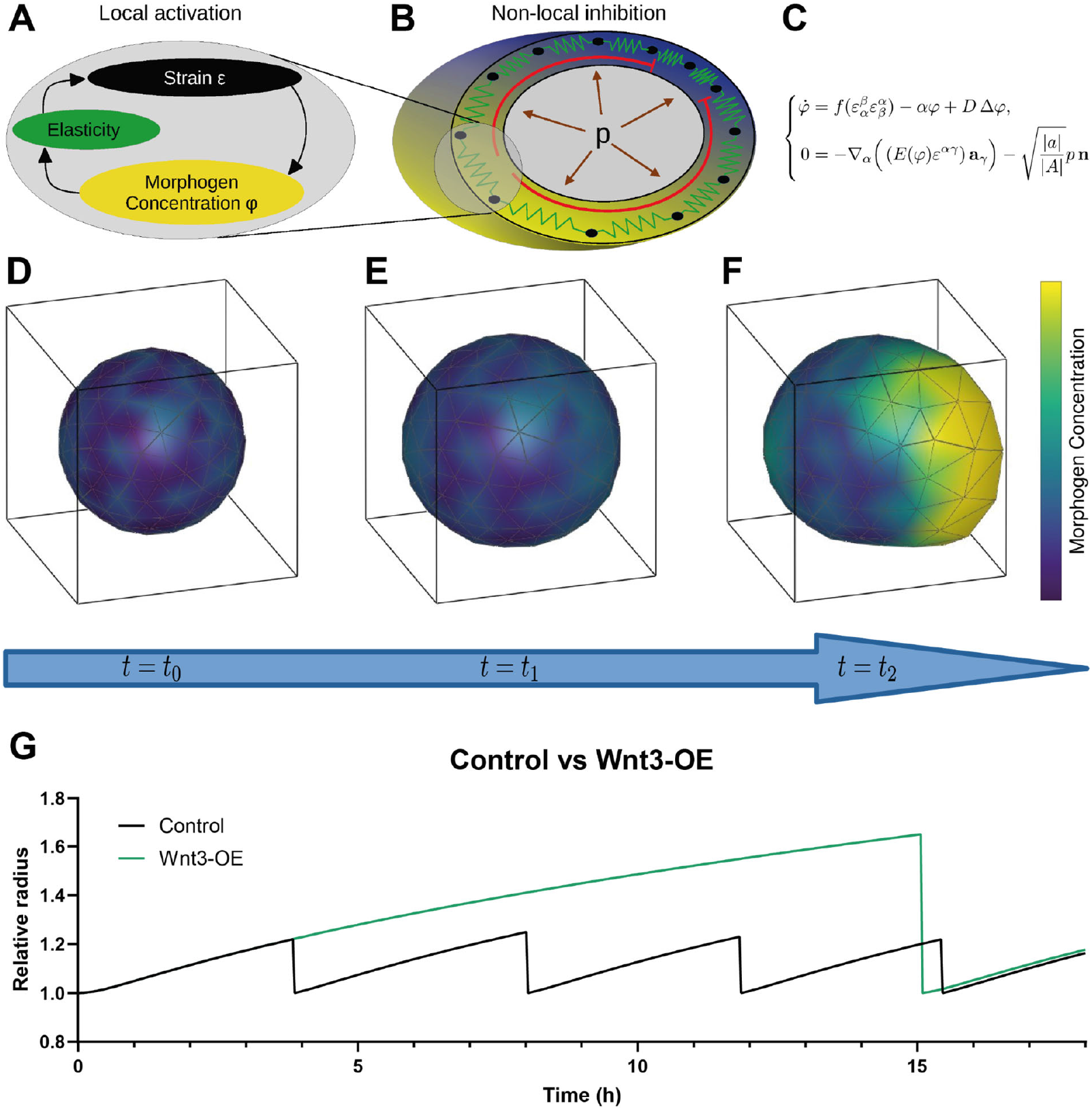
Mechanochemical model for symmetry breaking in *Hydra* spheroids. (**A**) Local activation through a positive feedback loop coupling strain and morphogen concentration. (**B**) Non-local inhibition mechanism driven by volume and cell count conservation. (**C**) Mathematical formulation of the mechanochemical model for pattern formation, combining a reaction-diffusion equation for morphogen concentration. 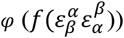 represents morphogen expression as a function of strain, *α* is the morphogen decay rate, *D* is the morphogen diffusion constant, Δ is the diffusion (Laplace-Beltrami) operator) and planar elasticity theory. The governing equation of planar elasticity of a shell couples the first Paola-Kirchhoff stress tensor (*E*(*φ*) represents Young modulus, *a*_y_ is the basis vector on deformed spheroid surface) with inner pressure p (**n** is outward unit normal direction on deformed surface, 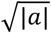 and 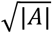 are surface measures on the deformed and undeformed surfaces). (**D**) Simulation of spontaneous pattern formation starting from spherical initial shape with randomly perturbed constant morphogen concentration. Over time, morphogen concentration and strain co-localize (**E**) leading to a local protrusion with high morphogen concentration (**F**). (**G**) Simulated trajectories showing the size dynamics of control and *Wnt3* overexpressing spheroids.

A lower-dimensional version of this model, representing the spheroid as a closed ring of fixed size, enables analytical examination. This approach allows us to delineate the pattern formation mechanism arising from the destabilization of the steady state. Linear stability analysis reveals that the growth rates of eigenmodes decrease as the number of peaks increases (Fig. S3, supplementary text). Consequently, patterns with a single peak in morphogen concentration tend to emerge, consistent with the observation that *Hydra* spheroids of various sizes generally develop a single body axis. This robustness constitutes a key difference from classical reaction-diffusion systems, such as the Gierer-Meinhardt model, which predict that the number of peaks scales with domain size.

### Mechanical stimulus localizes Wnt^+^ organizer

The comprehensive mechanochemical model for 3D spheroid patterning enables further evaluation of its explanatory power in response to experimental perturbations. One particularly intriguing observation, which cannot be fully explained by biochemical mechanisms alone, is the biasing of organizer location following mechanical stimulation. When suction is applied to a random site on the regenerating spheroid via pipette, the organizer consistently forms at a specific angle relative to the axis of suction (Sander, et al. 2020) (Fig. 3A-B, Movie S6).

**Figure 3:**
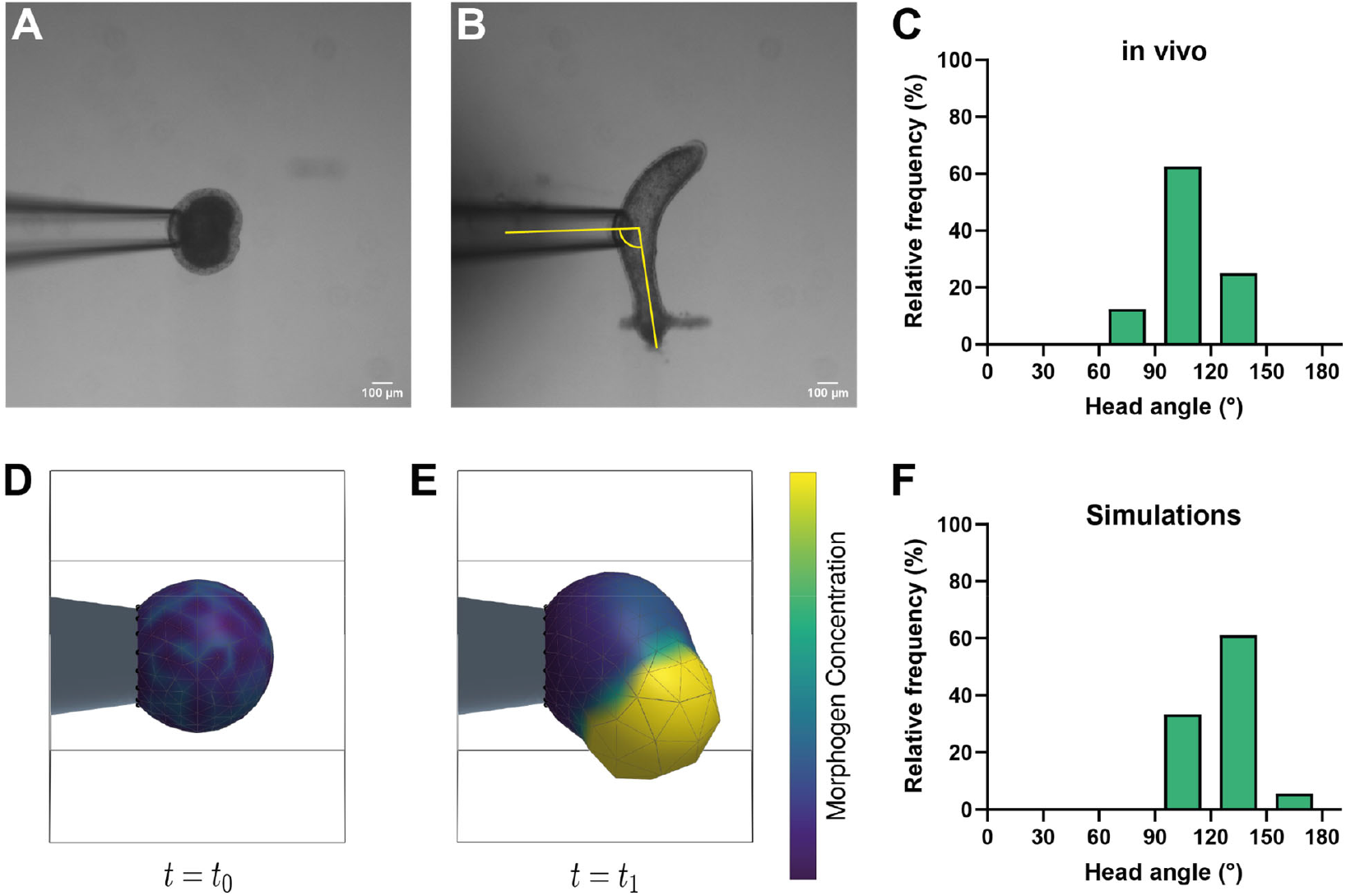
Pipette aspiration mechanically biases the angle of axis formation. Snapshots of *Hydra* spheroids regenerating while undergoing micropipette aspiration at the start (**A**) and end (**B**) of the regeneration process. (**C**) Distribution of axis formation angles in spheroids regenerating under pipette aspiration (n = 8). Angles were measured as indicated by the yellow lines in (B). The simulation of pattern formation under pipette aspiration begins at time *t*_*0*_, with a patch of nodes on the spheroid immobilized as if held in a pipette (**D**). By time *t*_*1*_, high morphogen expression localizes at a distinct angle from the immobilized nodes (**E**). (**F**) Distribution of axis formation angles in simulations mimicking pipette aspiration (n = 18). Angle distributions were compared using a Kolmogorov-Smirnov test: (C) vs. (F), *P* = 0.617; (C) vs. theoretically expected random distribution, *P* = 0.095. Scale bars, 100 μm.

Notably, the new head develops nearly perpendicular to the direction of the applied suction (Fig. 3C). We simulated this experiment by immobilizing a small patch of the spheroid, corresponding to the tissue trapped in the pipette, which does not stretch as much as the surrounding tissue during inflation (Fig. 3D). Consistent with experimental observations, the simulations reveal a high degree of tissue stretch around the rim of the pipette, while the tissue inside experiences far less deformation. Our simulations further show that the new organizer forms at an angle relative to the immobilized region, closely mirroring experimental results (Fig. 3E-F, Movie S7). This mechanochemical mechanism, therefore, provides insights into complex patterning phenomena that extend beyond those captured by traditional models.

### The most stretched area develops into an organizer

To further experimentally validate our model, we investigated the spatial relationship between the organizer and regions of maximal mechanical stretching, hypothesizing that these areas would coincide. Utilizing a transgenic line expressing GFP-tagged β-catenin under the control of the *β-catenin* promoter (Nakamura, et al. 2011), we visualized β-catenin localization through live imaging. Independent of Wnt signaling, β-catenin is found at cell-cell junctions, linking adherens junctions to the actin cytoskeleton. Upon Wnt activation, however, β-catenin translocates to the nucleus (Valenta, et al. 2012).This approach enables simultaneous observation of epithelial morphology and nuclear β-catenin localization. Our results confirmed that regions of maximum stretching, where cells flatten, coincided with elevated Wnt signaling activity (Fig. 4A-B). As predicted by our mechanochemical feedback model, the Wnt organizer consistently forms in the domain subjected to the greatest mechanical stimulation.

**Figure 4:**
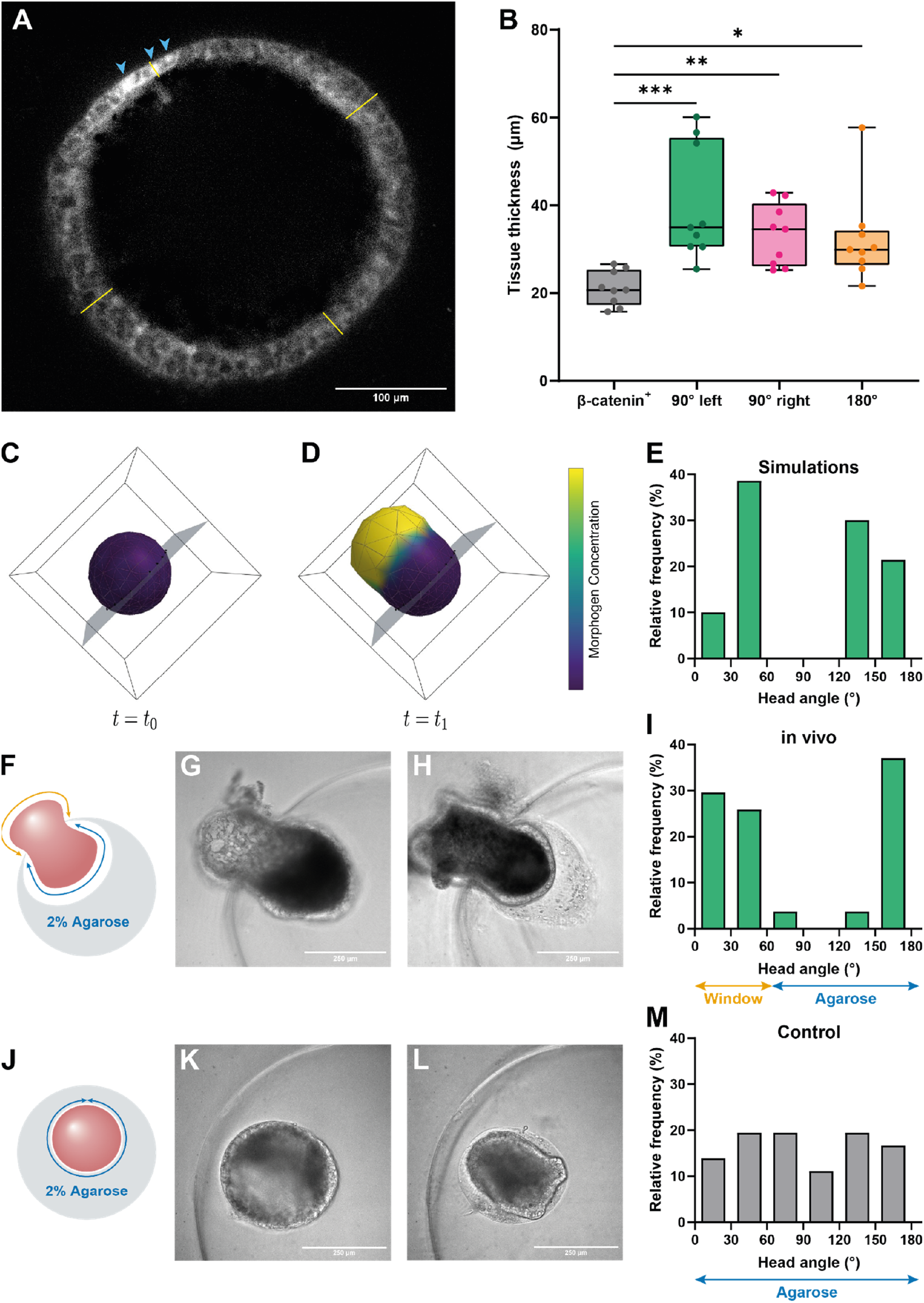
The location of the organizer correlates with regions of the highest tissue stretching and can be biased by mechanical confinement. (**A**) Optical section of a *β-catenin::GFP Hydra* spheroid at 19.2 hours, illustrating the variability in tissue thickness across its surface, with the organizer indicated by a localized patch of β-catenin^+^ nuclei (blue arrowheads). (**B**) Tissue thickness compared across the surface of regenerating spheroids at the first timepoint that clustered β-catenin^+^ nuclei are clearly visible. Tissue thickness was measured relative to the area containing β-catenin^+^ nuclei, as indicated by the yellow lines in (A) (*n* = 9). (**C**) Simulation of pattern formation under mechanical confinement where stretch is restricted in a single plane. (**D**) The final location of the protrusion associated with high levels of morphogen is significantly dependent on the immobilization plane. (**E**) Distribution of angles resulting from simulations of spheroids with confined stretch in a single plane (n = 70). (**F**) Schematic of the experimental setup mimicking mechanical confinement as shown in (C, D), where spheroids were partially embedded in 2% agarose, causing local confinement of stretch by the ring-shaped edge of the agarose upon inflation. Snapshots illustrate a confined, regenerating spheroid at maximum inflation (**G**) and after successful regeneration (**H**). (**I**) Distribution of angles of axis formation in spheroids experiencing local confinement in agarose (n = 27). (**J**) Schematic of the control setup with spheroids fully embedded in 2% agarose. Snapshots depict a fully embedded, regenerating spheroid at maximum inflation (**K**) and after successful regeneration (**L**). (**M**) Distribution of angles of axis formation in control spheroids fully embedded in 2% agarose (n = 36). Data in (B) were analyzed using a Kruskall-Wallis test, with Dunn’s multiple comparisons test for post-hoc analysis.. ^***^*P* < 0.001, ^**^*P* < 0.01, ^*^*P* < 0.05. Distributions were compared using a Kolmogorov-Smirnov test: (E) vs. (I): *P=*0.592; (I) vs. theoretically expected distribution: *P=*0.001; (M) vs. theoretically expected distribution: *P=*0.439. Scale bars, 100 μm (A) and 250 μm (H, I).

Building on these findings, which establish a clear link between the degree of stretching and cell fate toward becoming an organizer, we aimed to manipulate the spheroid in a way that would reliably influence organizer formation. Our model suggests that the organizer’s location can be influenced by altering the mechanical behavior of regenerating spheroids. Specifically, simulations indicate that introducing an expansion constraint, such as a ring, stimulates the formation of an organizer perpendicular to the plane of the constraint (Fig. 4C-E, Movie S8). To test this experimentally, we encased spheroids in 2% agarose gels, leaving a small open window through which the spheroid could expand into the medium (Fig. 4F). This setup allowed the agarose rim to contain the spheroid, enabling inflation-driven tissue stretching toward either the agarose cavity or the surrounding medium. The circular constraint of the window rim caused the spheroid to expand perpendicularly to the plane of the window (Fig. 4G-H, Movie S9), leading to the emergence of a new oral end at these sites (Fig. 4I). This specific pattern of head regeneration, aligned with the mechanical constraint, sharply contrasts with the random distribution observed when spheroids are encased in agarose without localized mechanical perturbations (Fig. 4J-M, Movie S10). The imposed asymmetry in tissue stretching during inflation biases the organizer’s localization. Thus, by manipulating the mechanical behavior of the tissue, we can direct the organizer’s emergence to a predetermined location.

## DISCUSSION

Our proposed mechanochemical mechanism for the emergence of a Wnt signaling center in regenerating *Hydra* tissue fragments presents an innovative alternative to the purely biochemical model suggested by Gierer and Meinhardt (Gierer and Meinhardt 1972). The seminal reaction-diffusion model, which has long been the dominant paradigm for pattern formation, relies on the co-dependent interaction of two diffusible factors with significantly different diffusion rates (Murray 2011). While pairs like Nodal and Lefty (Muller, et al. 2012) or BMP and Wnt (Raspopovic, et al. 2014) have been explored in other tissue patterning systems, no definitive pair has been identified in *Hydra*, despite extensive research. Although Wnt ligands remain crucial as activators, our mechanism eliminates the need for a second diffusible inhibitor by integrating mechanical components. In this case, mechanical forces do not simply modify the biochemical behavior of diffusible factors, as suggested in other models (Rauch and Millonas 2004; Zakharov and Dasbiswas 2021), but are essential elements of the patterning process itself.

Recent renewed interest in the mechanical properties of *Hydra* tissue has not only led to their detailed assessment (Perros, et al. 2024) but also highlighted their critical role in regeneration (Wang, et al. 2023). The recognized influence of mechanical events on *Hydra* regeneration has brought significant attention to the nematic order evident in the arrangement of supracellular actin fibers at the tissue level (Livshits, et al. 2017). These fibers align parallel to the main axis in epidermal epithelial cells, and their organization results in topological defects of +1 aster types at the animal’s extremities (Maroudas-Sacks, et al. 2021). A clear correlation exists between the localization of these topological defects and the Wnt^+^ organizer cells (Shani-Zerbib, et al. 2022). It has been recently proposed that they act as mechanical organizers of the morphogenetic events associated with regeneration (Ravichandran, et al. 2024). However, a comprehensive framework has been lacking. The causal relationship between actin cytoskeletal organization and Wnt signaling organizer emergence remains debated (Wang, et al. 2020) even though theoretical models have been proposed to resolve it where a link between the morphogen gradient and the orientation of filaments has been explored (Maroudas-Sacks, et al. 2024; Wang, et al. 2023). Importantly, differential stretching has been linked to distinct patterns of underlying actin filament organization (Bailles, et al. 2024; Maroudas-Sacks, et al. 2024). Ultimately, the mechanism we propose accommodates these observations, as variations in tissue stretching around topological defects influence the localization of *Wnt* expression.

Indeed, our approach can explain broader phenomena, such as the spontaneous *de novo* patterning of re-aggregated *Hydra* cells, where initial nematic order is absent during the emergence of Wnt organizers (Seybold, et al. 2016). The progressive emergence of a global nematic order has been documented and the fragmented organization of supracellular actin filament bundles is evident (Bailles, et al., 2024). Yet, under these conditions, the transmission of forces through supercellular actin cables is limited to smaller domains with interconnected actin networks. Ultimately, the spatial range of inhibition is restricted, enabling multiple organizer centers to emerge. It can thus be posited that the variabilities of the nematic order may act as a potential biasing mechanism, alongside a random distribution of chemical or mechanical factors.

In conclusion, we propose a mechanism that exploits mechanochemical variability within a closed epithelium to drive localized cell fate determination, responding to a base level of isotropic stress through asymmetric cellular responses. Similar phenomena are ubiquitous in luminal epithelial structures, including the otic vesicle (Mosaliganti, et al. 2019), liver canaliculi (Dasgupta, et al. 2018; Meyer, et al. 2020), and lung (Li, et al. 2018). This mechanism also influences early mammalian development, evidenced by documented cycles of inflation and deflation in blastocysts (Chan, et al. 2019). The proposed mathematical model may prove applicable to the patterning challenges faced by structures of this kind in general.

## MATERIALS & METHODS

### *Hydra* strains and culture conditions

All experiments were conducted using *Hydra vulgaris* (AEP strain). The specific transgenic lines used are noted for each experiment. *Hydra* were cultured in Volvic water at 18°C and fed freshly hatched *Artemia nauplii* three times per week. For the experiments, only animals without sexual organs and buds were selected. All procedures were carried out in compliance with ethical guidelines and Swiss/German national regulations for animal research.

### Spheroid preparation

The animals were transferred to *Hydra* medium (HM), composed of 1 mM CaCl_2_, 0.2 mM NaHCO_3_, 0.02 mM KCl, 0.02 mM MgCl_2_, and 0.2 mM Tris-HCl (pH 7.4), or to Volvic water for micropipette aspiration experiments. Spheroids were prepared following established protocols (Ferenc and Tsiairis 2022; Fütterer, et al. 2003). Briefly, animals were bisected at the mid-gastric region. From each half, a tissue ring was obtained by re-cutting the body axis. These rings were further divided into 2-4 rectangular pieces, depending on their size. Tissue fragments were allowed to fold into spheroids for 3-4 hours in dissociation medium (3.6 mM KCl, 6 mM CaCl_2_, 1.2 mM MgSO_4_, 6 mM sodium citrate, 6 mM sodium pyruvate, 4 mM glucose, and 12.5 mM N-tris(hydroxymethyl)methyl-2-aminoethanesulfonic acid, pH 6.9) at room temperature. Spheroids of the typical size (200-350 μm in diameter) and that had closed properly were selected for further experimentation.

### Live imaging of regenerating spheroids

Unless otherwise specified, regenerating spheroids were imaged in multichamber LabTek slides (Nunc, 155361) using individual agarose wells filled with HM. Agarose wells were prepared by coating the bottom of the chambers with a 2-to 3-mm layer of 1% agarose in HM. Holes were created using a 1 mL micropipette tip after the agarose solidified. The chambers were then filled with HM. For the experiments with alsterpaullone (AP) treatment, HM was first supplemented with 10 μg/mL Hoechst-33258. After 1.5-2 hours, the medium was replaced with HM containing either 0.2% DMSO (control) or 5 μM AP (Sigma, A4847) and samples were imaged immediately after. For experiments with genetic *Wnt3* overexpression, animals from the AEP *ecto pAct::eGFP* (Wittlieb, et al. 2006) and *pAct::Wnt3* lines (Ferenc, et al. 2021) were used.

Samples were imaged at 5- or 10-minute intervals for at least 24 hours at 18-21°C. For *Wnt3* overexpressing spheroids, imaging was conducted on a Zeiss LSM710 confocal laser scanning microscope with a 10x/0.45 Plan-Apochromat objective (Zeiss, 420640-9900-000). A single imaging plane per spheroid was acquired using bright-field and 571 nm laser illumination in a 512 × 512 format with a 2.2 μm pixel size. AP-treated samples were imaged using a Leica M205 FCA fluorescence stereo microscope with a 1.0x Plan-Apochromat objective (Leica, 10450028), manual zoom, and a Leica DFC480 camera. An overview image (1280 × 960, 5.4 μm pixel size) of all regenerating spheroids in a single field of view was acquired, and cropped frames containing individual spheroids were generated for later analysis.

### Micropipette aspiration and observation

Borosilicate glass capillaries (Science Products) with an outer diameter of 1 mm and an inner diameter of 0.58 mm (0.58×1.00×100 mm) were prepared using a P-97 pipette puller (Sutter Instruments). The pulled pipette ends were polished using a DeFonbrune-type microforge (Alkatel) to achieve blunt tips with an outer diameter of approximately 100 µm.

Experiments were conducted on an inverted Olympus IX51 microscope with a 4x objective (Olympus UPlanFl). Imaging was performed with a Hamamatsu C9100 Electron Multiplying Charge-Coupled Device (EMCCD) camera. The micropipette’s position was controlled using a XenoWorks electric micromanipulator (Sutter Instruments). The outer opening of the pipette was connected, via tubing, to a custom-made water-filled U-tube manometer.

Suction pressure (P_0_) was set using a syringe and maintained constant throughout the experiment. A P_0_ of 40 mm hydrostatic head (∼4×10^2^ Pa) allowed spheroids to be picked up from the dish bottom and moved to the observation area. Bright-field images were captured every 5 minutes in a 1000×1000 pixel format, minimizing illumination to avoid interference with regeneration.

### Long-term lightsheet imaging of regenerating spheroids

To capture tissue thickness across the entire surface of regenerating spheroids, we used a prototype of the LS2 Live lightsheet microscope (Viventis). Further details on this microscope can be found in (Moos, et al. 2024).

For each experiment, 6-8 spheroids of the *β-catenin::GFP* line (Nakamura, et al. 2011) were mounted in custom FEP chambers designed specifically for regenerating *Hydra* spheroids. Each chamber contained four pockets, each 0.5 mm in diameter in which a single spheroid could regenerate. The chambers were filled with 500 μL HM and any bubbles were removed from the pockets. Properly closed spheroids were transferred individually and directed into an empty pocket. The chambers were then fixed into a sample holder with room for four chambers, which was positioned on the microscope’s motorized stage.

To prevent overheating of the samples due to the lack of a cooling chamber on the microscope, several precautions were taken. The microscopy room was cooled using the room’s air conditioning system, and the outer chamber of the microscope was left partially open to allow for constant airflow between the cooled air of the room and the air inside the microscope.

Data were acquired using two 16x/0.8 long working distance (LWD) objectives (Nikon, MRP07220) and ORCA-Fusion sCMOS cameras (Hamamatsu, C14440-20UP). Illumination was provided by 488 nm and 561 nm lasers. Z stacks were taken for each spheroid by identifying the central plane and capturing 351 planes, spaced 2 μm apart, around this central plane. Each plane was recorded at a resolution of 2304 × 2304 pixels with an *xy* pixel size of 0.406 μm. Images were acquired every 10 minutes for at least 24 hours.

### Regenerating spheroids trapped in agarose

Spheroids from the *β-catenin::GFP* line were prepared and, after successful closure, partially embedded in 2% TopVision low melting point agarose (Thermo Scientific, R0801). The spheroids were submerged in melted agarose and gently sucked into a glass cylinder (Brand, 701904/701932) ensuring proper distribution. Subsequently, the spheroids were allowed to sink and make contact with the cylinder wall. After the agarose hardened, it was removed from the cylinder, and the portions containing a spheroid were separated. Spheroids sufficiently close to the agarose edge were selected, mounted in a glass-bottom chamber slide (Ibidi, 80427/80827), and covered with HM for imaging.

Imaging was performed using a Zeiss AxioObserver 7 microscope equipped with a CSU-W1 spinning disk confocal scanning unit (Yokogawa) featuring a 50 μm pinhole disk, a 20x/0.8 Plan-Apochromat objective (Zeiss, 440640-9903-000) and two Prime 95B sCMOS cameras (Photometrics). Visiview imaging software (Visitron) was used to control the system. For each spheroid, a single plane was acquired using bright-field and 488 nm laser illumination in parallel, at a 1200 × 1200 pixel resolution with a 0.55 μm pixel size. Imaging was conducted at 18-21°C, with frames captured every 10 minutes over a period of 67-69 hours.

### Quantification of oscillation parameters

The approach taken for quantification has been previously described (Ferenc and Tsiairis 2022). Briefly, bright-field images were systematically segmented using a Fiji macro (Schindelin, et al. 2012). The process began with a median filter (radius = 10 pixels), followed by automatic local thresholding (Phansalkar method, radius = 300 pixels, with r and k parameters set to 0). Any holes were filled, and object areas smaller than 50,000 μm^2^ that did not touch the image border were detected. The radius of each spheroid was calculated from the area, assuming a spherical shape. The radii were normalized per sample to either the first or lowest value. The normalized radius values and oscillation trajectories were imported in Microsoft Excel (Version 2406 Build 16.0.17726.20078) to calculate the maximum radius, average oscillation period, and average slope of phase I oscillations. The slope was determined using the trendline function in Excel, focusing solely on the inflating segments of the oscillations. For all radius values relevant to these parameters, the images were double-checked to prevent skewing of the data due to mis-thresholding.

Samples that died during imaging were excluded from analysis. Additionally, considering the time it takes for AP treatment to take effect, spheroids treated with AP that started displaying AP-driven behavior (e.g., larger oscillations) only when less than one full oscillation period (median = 745 minutes, calculated from AP-treated samples) was left, were also excluded from the analysis.

### Quantification of the angle of head formation

For regenerating spheroids under pipette aspiration, the head angle was determined as the first tentacle(s) emerged. The angular position of the future mouth relative to the pipette axis was measured using the Fiji angle tool. Due to *Hydra*’s movement, there was some uncertainty in the angle, so we estimated an angle rounded to the nearest 10 degrees based on several measurements at different time points for the same *Hydra*.

Although agarose-embedded spheroids were selected for imaging based on their proximity to the edge of the agarose, many were still fully embedded. Spheroids with any visible “window” (an opening to the medium) were considered, while fully embedded spheroids served as controls. Spheroids that died, failed to regenerate, or escaped the agarose during regeneration were excluded. Additionally, spheroids that rotated before tentacle buds appeared or regenerated outside the *xy* plane, were also omitted from analysis.

The head formation angle was measured at the first time that tentacle buds became visible using Fiji’s angle tool. For partially embedded animals, the head formation angle was measured relative to the center of the “window” in the agarose. For fully embedded spheroids, the angle was measured relative to the thinnest area of the agarose.

For simulated spheroids under pipette aspiration, the head angle was determined by assessing the maximum volume state immediately before the final simulated rupture. All nodes with a morphogen concentration higher than 0.75 were selected as part of the pattern cluster. The mean position of these nodes was calculated in cartesian space, and the angle between this center and the pipette axis was measured.

When simulating partial confinement by agarose the head angle was assessed at the maximum volume state immediately before the final simulated rupture. All nodes with a morphogen concentration higher than 0.75 were selected as part of the pattern cluster. The mean position of these nodes was calculated in cartesian space, and the angle between this center and the center of the “window” in the agarose was measured. In cases of near-symmetrical confinement when high-morphogen patches would appear in both halves of the spheroid, the patch with larger mass of morphogen would be measured.

### Measuring tissue thickness in regenerating spheroids

Before quantifying tissue thickness, the lightsheet datasets were extensively processed. For fusion of the data acquired by the individual detection objectives, we adapted previously described Python scripts (Moos, et al. 2024). In summary, image stacks obtained from the two objectives were fused using a sigmoidal function, centered at the plane where the image quality transitions between the two cameras. This plane was determined manually for each sample and was kept constant across all time points. The fusion process generated two image stacks per sample for each time point, corresponding to each illumination. To minimize autofluorescence in the GFP channel, the image stack from the 561 nm laser was multiplied by a factor of 2.2 and subtracted from the 488 nm image stack in Fiji.

Tissue thickness was measured for all samples that successfully regenerated within 66 hours. The first time point and individual z-plane where clustering of β-catenin^+^ nuclei was clearly visible were identified. Tissue thickness was measured using the membranal β-catenin signal in Fiji. To create a cleaner edge of the tissue, a median filter (radius = 5 pixels) was applied. Measurements began in the center of the area containing β-catenin^+^ nuclei. Locations 90° on either side and 180° opposite this area were also determined and measured.

### Statistical analysis

For the statistical comparison of oscillation dynamics and tissue thickness, GraphPad Prism 10 was utilized. To compare two samples with no significant variance difference, an unpaired, two-sided t-test was applied. Alternatively, a two-tailed Mann-Whitney test was used. For comparisons involving more than two samples, a Kruskal-Wallis test was conducted, followed by Dunn’s multiple comparisons test for *post-hoc* analysis.

For the comparison of angle distributions, the open-source software R (R Core Team 2024) was employed. Two variants of the Kolmogorov-Smirnov test were performed. First, to compare simulated and experimental head angle distributions, a standard two-sided Kolmogorov-Smirnov test using the function *ks*.*test()* was utilized. Second, to compare experimental results to a theoretically expected distribution, we assume random head formation for the latter. Given the area effects on the sphere (e.g., the area near the poles (0° and 180°) is sparser than the equatorial region (90°)), the theoretically expected cumulative distribution function was calculated as (1−cos(x))/2 (where angle x is given in radians). The function *ks*.*test()* (two-sided) was again used to compare the experimental data against this cumulative distribution function.

All plots were generated using GraphPad Prism 10. In all box plots, the box extends from the 25th to the 75th percentiles, with the central line representing the median. Whiskers extend to the minimum and maximum values, and all values are displayed as individual points. For histograms, bins of 30° were used, with bin boundaries indicated on the x-axis of each histogram.

## ACKNOWLEDGEMENTS

We would like to thank Jacqueline Ferralli for her technical support and assistance with *Hydra* upkeep throughout the project. We are grateful to Franziska Moos, Simon Suppinger, and Gustavo Quintas Glasner de Medeiros for providing access to their light sheet microscope and for their invaluable guidance. We also extend our appreciation to the Facility for Advanced Imaging and Microscopy at FMI, particularly Laure Plantard, for their support, and to Sara Raquel Roig Merino from the Imaging Core Facility of the Biozentrum for her generous introduction to light sheet microscopy. Special thanks to the Tsiairis, Großhans, Liberali, and Turco labs, Anaïs Bailles, and Pavel Tomancak for insightful discussions, and to Helge Großhans for critically reviewing the manuscript.

## Notes

### Competing Interest Statement

The authors have declared no competing interest.

